# Endocytosis and Compartmentalized Intracellular Signaling of the Prostaglandin Receptor EP4 Mediate Pain

**DOI:** 10.64898/2026.09.18.752773

**Authors:** Badr Sokrat, Raquel Tonello, Paz Duran, Naomi Barrett, Yatendra Mulpuri, Brian L. Schmidt, Dane D. Jensen, Francesco De Logu, Romina Nassini, Pierangelo Geppetti, Nigel W. Bunnett

## Abstract

Prostaglandin E_2_ (PGE_2_) is recognized as a major mediator of inflammatory pain. However, the intracellular signaling mechanisms by which PGE_2_ mediates pain remain unclear. Here, we show that pain-like responses evoked by PGE_2_ in dorsal root ganglion (DRG) nociceptors are mediated by internalization and compartmentalized signaling of the EP4 receptor. The EP4-selective agonist L-902,688 induced immediate nocifensive behavior and prolonged mechanical allodynia in mice and sensitized isolated DRG nociceptors. Pharmacological inhibition of clathrin- and dynamin-mediated endocytosis (using pitstop2 and dyngo4a) and siRNA knockdown of *Dnm1* prevented EP4-induced nociception and sensitization, implicating receptor trafficking in pain signaling. Using genetically-encoded biosensors, we monitored EP4 trafficking and downstream signaling within subcellular compartments of HEK293 cells. PGE_2_ stimulated dynamin- and β-arrestin-dependent EP4 trafficking from the plasma membrane to early, late and recycling endosomes and Golgi apparatus, and mobilized intracellular EP4 pools from the endoplasmic reticulum. PGE_2_ induced the assembly of EP4, G proteins and β-arrestin signaling complexes in endosomes and the Golgi apparatus. Inhibition of EP4 endocytosis suppressed intracellular cAMP production and ERK activation. Our findings reveal that EP4-mediated nociceptive signaling originates from intracellular compartments and suggest that modulating EP4 internalization and subcellular signaling dynamics could offer a novel strategy for inflammatory pain management.

**Teaser:** EP4 receptor endocytosis and compartmentalized intracellular signaling contribute to nociception and pain sensitization.

## Introduction

Chronic pain is a debilitating condition that afflicts 20-45% of individuals worldwide, yet remains inadequately managed for many patients (*1, 2*). Non-steroidal anti-inflammatory drugs (NSAIDs) are commonly used to treat pain. By inhibiting cyclooxygenase (COX) enzymes, NSAIDs block the synthesis of prostaglandins (PGs), the key mediators of inflammation and pain (*3, 4*). While NSAIDs can relieve acute pain, their continuous use causes adverse effects on the gastrointestinal, cardiovascular, hepatic and renal systems due to suppression of the protective effects of PGs. These life-threatening events restrict the long-term use of NSAIDs and highlight the need for a better understanding of how PGs cause pain (*5–7*).

PGE_2_ is a key inflammatory mediator that plays a central role in the development and maintenance of pain hypersensitivity by activating EP1-4 receptors. Of the four EP subtypes, EP4 has been linked to PGE_2_-induced activation and sensitization of nociceptors (*8*). EP4 is a G protein-coupled receptor (GPCR) that couples to Gαs proteins to activate adenylyl cyclase and thereby stimulate cyclic adenosine monophosphate (cAMP) production and protein kinase A (PKA) activation (*9, 10*). EP4 can also couple to inhibitory Gαi proteins (*11*), indicating that this receptor can engage multiple downstream pathways.

GPCRs are the largest family of transmembrane receptors in humans and constitute a major focus for drug development, with approximately 34% of FDA-approved drugs targeting GPCRs (*12, 13*). GPCRs are conventionally considered to signal primarily from the plasma membrane by binding extracellular agonists and coupling to intracellular heterotrimeric G proteins. The four main families of G proteins regulate different effectors, including adenylyl cyclase (Gαs, Gαi), phospholipase C (Gαq), RhoGEF (Gα12), ion channels and phosphoinositol-3-kinase (Gβγ), and thereby transduce distinct signaling pathways (*14, 15*). GPCR activation at the plasma membrane is tightly regulated. GPCR kinases phosphorylate activated GPCRs, which promotes the recruitment of β-arrestins. β-arrestins uncouple GPCRs from G proteins, which desensitizes signaling, and couple GPCRs to clathrin and adaptor protein 2, leading to dynamin-mediated endocytosis (*16, 17*). Desensitization and endocytosis terminate plasma membrane signaling. However, accumulating evidence indicates that many GPCRs can continue to signal from intracellular compartments (*18–20*). Activated GPCRs remain associated with Gα and β-arrestin isoforms in early endosomes, leading to sustained second messenger formation and kinase activation (*21*). Notably, several GPCRs internalize and continue to signal from endosomes to regulate pain, including protease-activated receptor-2 (PAR_2_) (*22*), calcitonin-like receptor (CLR) (*23*) and neurokinin 1 receptor (NK_1_R) (*24*).

We recently provided pharmacological and genetic evidence that PGs released during inflammation activate EP2 and EP4 to cause pain by mechanistically distinct pathways in mice (*25*). Pain-like behaviors induced by EP2 activation are mediated by Schwann cells and occur independently of the inflammatory response. EP2-driven cAMP/PKA signaling in Schwann cells is confined to the plasma membrane, implicating a spatially confined signaling mechanism. In contrast, EP4 activation elicits an acute nociceptive response through direct actions on nociceptors in dorsal root ganglia (DRG). EP4 undergoes ligand-induced internalization in response to PGE_2_, suggesting a fundamentally different mode of signal regulation (*25*). These findings raise critical questions about the role of EP4 trafficking and intracellular signaling in nociceptor sensitization and pain, suggesting the need for further investigation into the mechanisms and functional consequences of EP4 endocytosis in the context of pain.

We hypothesized that EP4 endocytosis and intracellular signaling contribute to nociception and pain sensitization. Our results show that activated EP4 undergoes β-arrestin and dynamin-mediated endocytosis and that inhibitors of endocytosis block EP4-induced nociceptive behavior and sensitization of nociceptors, providing a link between receptor endocytosis and pain. We determined the contribution of dynamin and β-arrestin to EP4 intracellular trafficking and signaling as well as the role of EP4 endocytosis in second messenger production and kinase activation. Our findings reveal a novel mechanism of sustained pain signaling that could be selectively targeted to achieve analgesic effects while avoiding the adverse effects associated with current pain treatments.

## Results

### Endocytosis mediates EP4-induced nociceptive behavior

We recently reported that the intraplantar (i.pl.) injection of the EP4-selective agonist, L-902,688, in the mouse hind paw produced immediate and transient (10-20 min) licking and lifting of the injected paw, indicative of acute nocifensive responses, and reduced withdrawal thresholds to stimulation of the ipsilateral paw with von Frey filaments for at least 4 hours, consistent with sustained mechanical allodynia (*25*). These pain-like responses were attenuated by the selective EP4 antagonist, BGC20-1531. The primary role of DRG neurons in mediating nocifensive responses and mechanical allodynia to the EP4-selective agonist, L-902,688, was revealed by the reduction of both nocifensive responses and mechanical allodynia observed in mice with selective downregulation of EP4 expression produced via the injection of *Adv^Cre^*drivers to express short hairpin RNA (shRNA) selectively in DRG neurons of *Adv^Cre^*mice (*25*).

To determine the contribution of endocytosis to EP4-induced nociceptive behavior in mice, we investigated the effects of inhibitors of clathrin and dynamin on the immediate nocifensive responses and sustained mechanical allodynia to the EP4-selective agonist L-902,688. The clathrin inhibitor pitstop2 (500 pmol/10 µl), the dynamin inhibitor dyngo4a (500 pmol/10 µl) or vehicle (PBS, 10 µl, control) were administered by i.pl. injection into the left hindpaw 30 min before injection of the EP4 agonist, L-902,688 (5 nmol/10 µl) into the same site. In vehicle-treated mice, L-902,688 stimulated acute nocifensive responses and reduced withdrawal thresholds to stimulation of the ipsilateral paw with von Frey filaments for at least 4 hours, consistent with sustained mechanical allodynia. Pitstop2 or dyngo4a completely prevented L-902,688-evoked nocifensive responses and mechanical allodynia (**Fig. 1A, B**), providing evidence for a role for endocytosis in EP4-mediated nociception.

**Figure 1:**
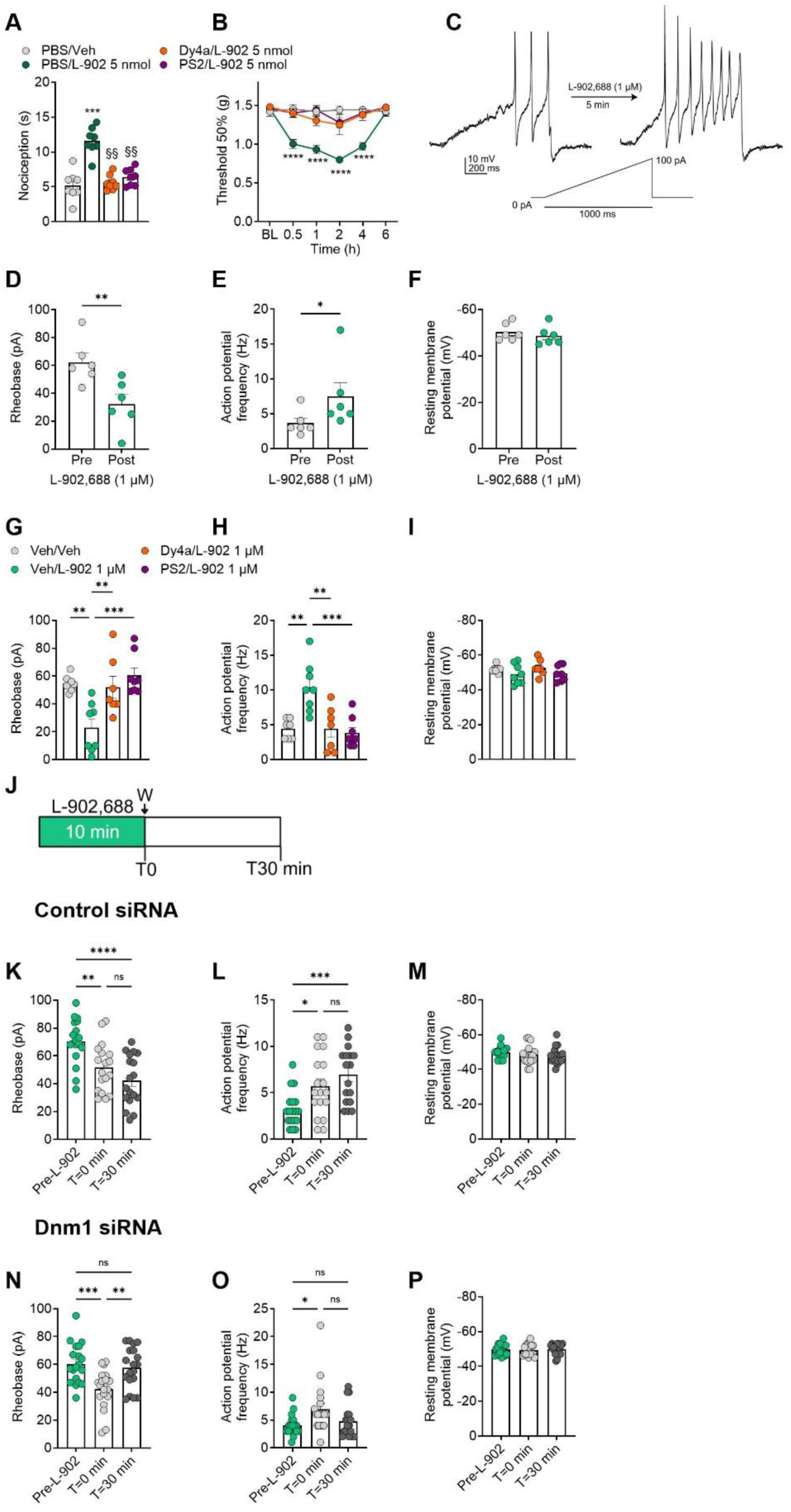
EP4 receptor activation mediates PGE_2_-evoked nociception and induces endocytosis-dependent sensitization of nociceptors. **A, B)** Effect of pretreatment with endocytic inhibitors, dyngo4a (Dy4a, 500 pmol/ 10µl, dynamin inhibitor) or pitstop2 (PS2, 500 pmol/ 10µl, clathrin inhibitor) on nocifensive behavior **(A)** or mechanical allodynia **(B)** induced by intraplantar (i.pl.) injection of L-902,688 (L-902, 5 nmol/ 10µl) or vehicle (Veh, 10µl) in mice (n=8 mice per group). **C)** Representative traces of action potentials in DRG neurons before and after application of the EP4 agonist L-902,688 (1 µM, 5 min). **D-F)** Rheobase **(D)**, action potential firing frequency **(E)** and resting membrane potential **(F)** pre- and post-L-902,688 stimulation (n=6). **G-I)** Effect of pretreatment with endocytic inhibitors, dyngo4a (Dy4a, 30 µM, dynamin inhibitor, n=7), pitstop2 (PS2, 15 µM, clathrin inhibitor, n=8) or vehicle (Veh, n=7), on L-902,688-induced neuronal hyperexcitability measured 30 min post-stimulation. Rheobase **(G)**, firing frequency **(H)**, and resting membrane potential **(I)** were measured. **J)** Experimental timeline for electrophysiology experiments 2 days after the intrathecal administration of control siRNA **(K-M)** or dynamin 1 (Dnm1) siRNA (N-P) in mice. DRG neurons were challenged with L-902,688 and washed. Rheobase, action potential firing frequency and resting membrane potential were measured at T=0 and T=30 min after L-902,688 challenge (n=18-19). Data are represented as mean ± SEM. Paired *t*-test, 1-way or 2-way ANOVA with Bonferroni correction or Tukey’s multiple comparison test. *P<0.05, **P<0.01, ***P<0.001, ****P<0.0001 vs. Veh, Pre-L-902, Veh/Veh or Veh/L-902; ^§§^P<0.01 vs. PBS/L-902.

### Endocytosis mediates EP4-induced sensitization of nociceptors

To assess the impact of EP4 activation on DRG neuron excitability, we made patch clamp recordings from isolated mouse DRG neurons. Neurons were challenged with L-902,688 (1 µM, 5 min) and rheobase (minimum current to fire action potentials) and action potential frequency were measured. L-902,688 significantly decreased the rheobase (pre: 62.3 ± 6.5 pA; post: 32.0 ± 7.1 pA, n=6) and increased action potential firing frequency (pre: 3.6 ± 0.7 Hz; post: 7.5 ± 1.9 Hz; *P*<0.01, paired *t*-test), without altering the resting membrane potential (**Fig. 1C-F**).

To investigate whether endocytosis contributes to EP4-induced neuronal hyperexcitability, we pretreated DRG neurons with inhibitors of clathrin- and dynamin-mediated endocytosis. Neurons were preincubated with dyngo-4a (30 µM) or pitstop-2 (15 µM) for 10 min, challenged with L-902,688 (1 µM) excitability was measured 30 min post-stimulation. Compared to vehicle-treated neurons (rheobase: 23.0 ± 6.2 pA; frequency: 4.4 ± 0.5 Hz; n=8), dyngo-4a and pitstop-2 pretreated neurons exhibited significantly increased rheobase (52.0 ± 7.9 pA; n=7 and 60.7 ± 5.1 pA; n=8, respectively) and reduced firing frequency (4.4 ± 1.2 Hz and 3.8 ± 0.7 Hz, respectively; *P*<0.05 vs. vehicle, one-way ANOVA with Tukey’s test) (**Fig. 1G-I**).

To determine the specific role of dynamin 1, one of the most abundantly expressed dynamin isoforms in DRG neurons(*26*), in mediating endosomal EP4 signaling related to sustained nociceptor hyperexcitability, we evaluated the effects of L-902,688 on DRG neurons from mice treated intrathecally with control or *Dnm1* siRNA. DRG were collected two days after siRNA injection. We have previously shown that *Dnm1* siRNA knocksdown *Dnm1* mRNA in DRG neurons by ∼70% after two days (*27*). DRG neurons were preincubated with L-902,688 (1 µM, 10 min), washed and rheobase and action potential firing frequency were measured 0 or 30 minutes later (**Fig. 1J**). In neurons from control siRNA-treated mice, L-902,688 induced significant hyperexcitability at both time points (0 min: rheobase 51.6 ± 4.0 pA, frequency 5.7 ± 0.7 Hz; 30 min: rheobase 42.3 ± 4.3 pA, frequency 6.9 ± 0.7 Hz; n=18, *P*<0.05 vs. baseline, one-way ANOVA) (**Fig. 1K, L**). In contrast, dynamin 1 knockdown selectively reversed the sustained effects measured at 30 min (rheobase: 57.6 ± 3.3 pA; frequency: 4.7 ± 0.6 Hz; n=19, *P*>0.05 vs. control), while the acute response at 0 min remained intact (rheobase: 42.2 ± 3.2 pA; frequency: 6.9 ± 1.0 Hz; n=19, *P*<0.05 vs. control) (**Fig. 1N, O**). Resting membrane potential remained unchanged across all conditions and time points (**Fig. 1M, P**).

Together, these results demonstrate that EP4 activation induces sustained hyperexcitability of nociceptors and persistent mechanical allodynia through an endocytosis-dependent mechanism, implicating endosomal EP4 signaling as a critical driver of PGE_2_-mediated pain.

### Dynamin and β-arrestin mediate agonist-induced EP4 endocytosis and intracellular trafficking

To elucidate the mechanisms by which EP4 intracellular signaling may contribute to pain, we first characterized EP4 trafficking between different intracellular compartments. We used enhanced bystander Bioluminescence Resonance Energy Transfer (ebBRET) biosensors (*28*) to analyze the kinetics of the movement of EP4 between the plasma membrane and intracellular compartments after agonist stimulation. We coexpressed in parental HEK293 cells or β-arrestin 1/2 knockout (βarrKO) HEK293 cells, EP4 fused at the C-terminus to the BRET donor *Renilla* luciferase (Rluc8), and cellular compartment markers coupled to the BRET acceptor *Renilla* green fluorescent protein (rGFP or tandem rGFP) to track receptor localization in specific organelles (*29, 30*) (**Fig. 2A**). This approach enables high sensitivity and robust detection of protein trafficking due to the natural affinity and optimal energy transfer between the BRET donor and acceptor (*31*).

**Figure 2:**
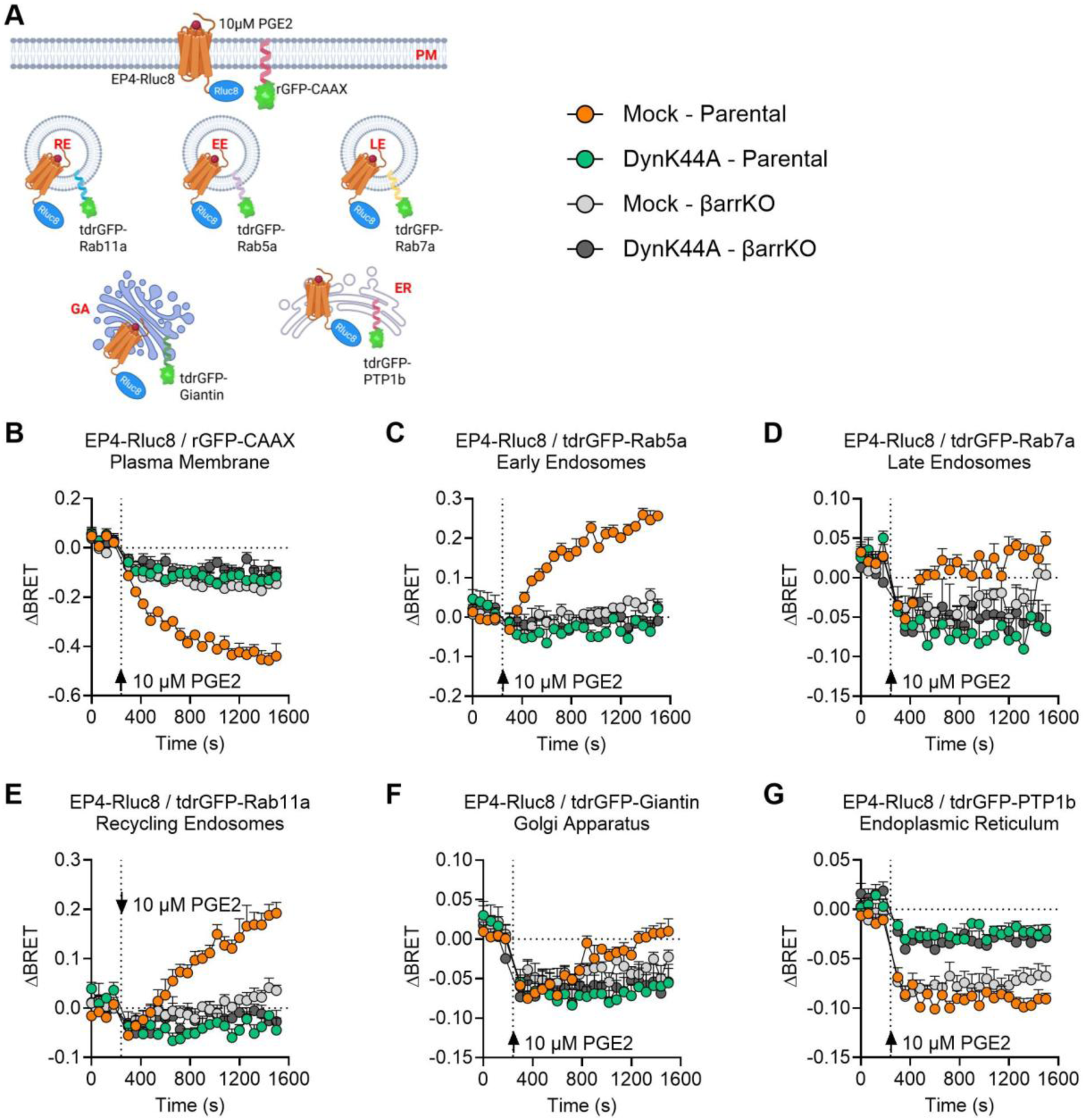
EP4 receptor trafficking monitored by enhanced bystander BRET (ebBRET). **A)** Illustration of the ebBRET biosensors used to measure EP4 receptor trafficking in subcellular compartments. **B-G)** ebBRET kinetic assay of PGE_2_-induced (10μM) trafficking of EP4-Rluc8 at **B)** the plasma membrane (rGFP-CAAX), **C)** early endosomes (tdrGFP-Rab5a), **D)** late endosomes (tdrGFP-Rab7a), **E)** recycling endosomes (tdrGFP-Rab11a), **F)** Golgi apparatus (tdrGFP-Giantin), and **G)** endoplasmic reticulum (tdrGFP-PTP1b) in parental HEK293 cells or β-arrestin1/2 KO cells with or without DynK44A expression. Data are represented as the mean ± SEM (n=5).

In parental HEK293 cells, PGE_2_ (10 μM) decreased the BRET signal between EP4-Rluc8 and the plasma membrane marker rGFP-CAAX, indicating receptor internalization (**Fig. 2B**). Expression of DynK44A, a dominant-negative mutant of dynamin, markedly inhibited the agonist-stimulated decrease in BRET at the plasma membrane. Agonist-stimulated EP4 internalization was significantly reduced in βarrKO cells and expression of DynK44A in these cells showed no further inhibitory effect (**Fig. 2B**). In parallel, we monitored the EP4 trafficking to early endosomes by measuring BRET between EP4-Rluc8 and the early endosome marker tdrGFP-Rab5a (**Fig. 2C**). In parental cells, PGE_2_ progressively increased the BRET signal between EP4-Rluc8 and tdrGFP-Rab5a, denoting endocytosis (**Fig. 2C**). Consistent with our observations at the plasma membrane, expression of DynK44A or absence of β-arrestin 1/2 completely inhibited EP4 endocytosis. These findings indicate that EP4 undergoes dynamin- and β-arrestin-dependent endocytosis.

To investigate the post-endocytic fate of EP4, we used ebBRET to monitor its proximity to markers of late endosomes, recycling endosomes, the Golgi apparatus, and the endoplasmic reticulum (ER). In parental HEK293 cells, PGE_2_ induced a small but significant increase in BRET signal between EP4-Rluc8 and the late endosome marker tdrGFP-Rab7a, suggesting that a fraction of endocytosed EP4 is sorted to late endosomes (**Fig. 2D**). Expression of DynK44A and absence of β-arrestin 1/2 abolished EP4 trafficking to late endosomes. PGE_2_ stimulated a robust increase in BRET signal between EP4-Rluc8 and the recycling endosome marker tdrGFP-Rab11a, consistent with predominant EP4 trafficking to a recycling pathway, which was similarly inhibited by DynK44A expression and in absence of β-arrestin 1/2 (**Fig. 2E**).

We similarly evaluated EP4 trafficking to the Golgi apparatus by measuring BRET between EP4-Rluc8 and the cis-Golgi marker tdrGFP-Giantin, and observed that PGE_2_ stimulated a delayed increase in BRET signal (**Fig. 2F**). Expression of DynK44A and absence of β-arrestin 1/2 inhibited this response, suggesting that EP4 undergoes endosome-to-Golgi retrograde transport (**Fig. 2F**). Finally, we monitored EP4 localization in the ER by measuring BRET between EP4-Rluc8 and the ER marker tdrGFP-PTP1b, and observed that PGE_2_ stimulated a sustained decrease in BRET signal indicating EP4 trafficking away from the ER, presumably to repopulate the plasma membrane with freshly synthesized receptors (**Fig. 2G**). While DynK44A expression inhibited PGE_2_-stimulated movement of EP4 from the ER, depletion of β-arrestin 1/2 had no effect. This suggests that the trafficking of the EP4 receptor away from the ER is dynamin-dependent and β-arrestin-independent.

Taken together, these results suggest that EP4 leaves the plasma membrane and traffics through the endosomes and Golgi apparatus using dynamin- and β-arrestin-mediated pathways while ER exit only requires dynamin.

### EP4 remains active in intracellular compartments

To determine the contribution of β-arrestin isoforms to EP4 trafficking, we monitored PGE_2_-induced BRET between β-arrestin 1 and β-arrestin 2 fused to RlucII (βarr1-RlucII, βarr2-RlucII) and rGFP-CAAX (plasma membrane), tdrGFP-Rab5a (early endosomes), tdrGFP-Rab7a (late endosomes), tdrGFP-Rab11a, tdrGFP-Giantin (cis-Golgi) or tdrGFP-PTP1b (ER) (**Fig. 3A**). PGE_2_ rapidly stimulated recruitment of both β-arrestin isoforms to the plasma membrane, with β-arrestin 2 displaying a transient peak before declining, while β-arrestin 1 recruitment was more sustained (**Fig. 3B**). At the early, late, and recycling endosomes, β-arrestin 1 and β-arrestin 2 exhibited a more sustained recruitment but at a slower rate compared to the plasma membrane (**Fig. 3C, D, E**). This is a consistent pattern of class B GPCRs, which form stable complexes with β-arrestin in endosomal compartments. We also observed sustained recruitment of β-arrestin 1 and β-arrestin 2 to the Golgi apparatus, suggesting a potential role for β-arrestin in the endosome-to-Golgi retrograde trafficking of EP4 or in scaffolding signaling complexes within this organelle (**Fig. 3F**). In contrast, β-arrestin 1 and β-arrestin 2 recruitment to the ER was transient potentially due to the departure of the EP4 receptor from this compartment (**Fig. 3G**).

**Figure 3:**
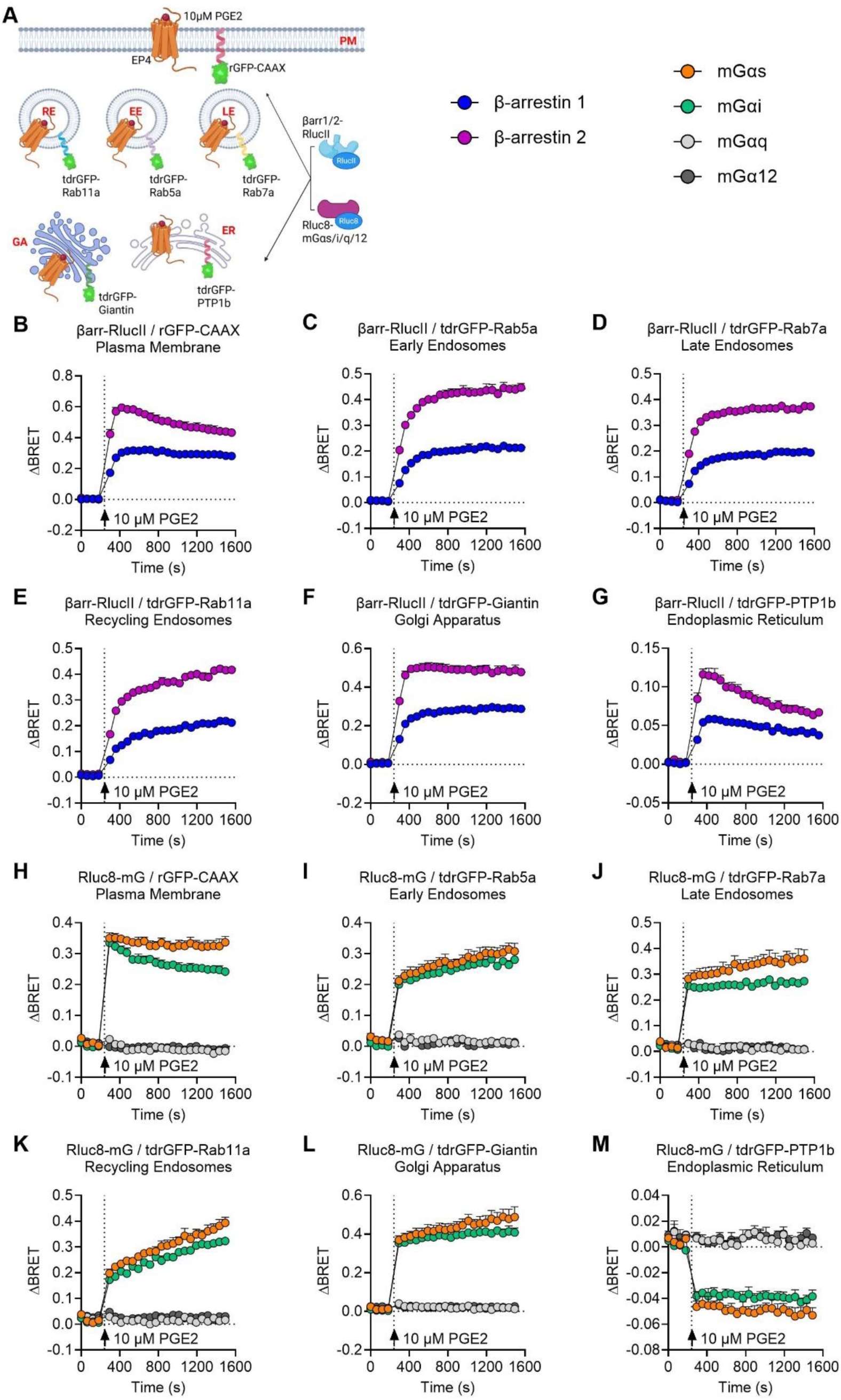
mGα proteins and β-arrestin1/2 recruitment to active compartmentalized EP4 receptor. **A)** Illustration of the ebBRET biosensors used to measure mGα protein and β-arrestin1/2 recruitment to subcellular compartments. **B-G)** PGE_2_-induced (10 μM) β-arrestin1/2 recruitment is monitored by ebBRET in HEK293 cells cotransfected with β-arrestin1/2-RlucII, EP4 receptor and ebBRET markers of **B)** the plasma membrane (rGFP-CAAX), **C)** early endosomes (tdrGFP-Rab5a), **D)** late endosomes (tdrGFP-Rab7a), **E)** recycling endosomes (tdrGFP-Rab11a), **F)** Golgi apparatus (tdrGFP-Giantin), and **G)** endoplasmic reticulum (tdrGFP-PTP1b). **H-M)** PGE_2_-induced (10μM) mGαs, mGαi, mGαq, and mGα12 recruitment is monitored by ebBRET in HEK293 cells cotransfected with Rluc8-mGα, EP4 receptor and ebBRET markers of **H)** the plasma membrane (rGFP-CAAX), **I)** early endosomes (tdrGFP-Rab5a), **J)** late endosomes (tdrGFP-Rab7a), **K)** recycling endosomes (tdrGFP-Rab11a), **L)** Golgi apparatus (tdrGFP-Giantin), and **M)** endoplasmic reticulum (tdrGFP-PTP1b). Data are represented as the mean ± SEM (n=4-5).

To assess the activation state of EP4 in different subcellular compartments, we measured PGE_2_-induced recruitment of mini Gα (mGα) protein isoforms fused to Rluc8 using ebBRET (**Fig. 3A**). mGα proteins are engineered versions of G proteins with a truncated N-terminal and α-helical domain that selectively recognize active conformations of GPCRs (*32, 33*). We comprehensively analyzed the EP4 activation state by measuring the recruitment of the four Gα protein families, mGαs, mGαi, mGαq and mGα12. PGE_2_ stimulated the rapid recruitment of mGαs and mGαi but not Gαq and Gα12 to the plasma membrane, consistent with previously observed EP4 coupling to mGαs and mGαi (**Fig. 3H**). PGE_2_ stimulated a gradually increasing and sustained recruitment of mαGs and mGαi to early endosomes, suggesting that EP4 remains active and capable of Gα protein coupling after endocytosis (**Fig. 3I**). A similar pattern was observed at the late and recycling endosomes, where PGE_2_ stimulated robust recruitment of mGαs and mGαi. We also detected that PGE_2_ stimulated mGαs and mGαi recruitment to the Golgi apparatus, suggesting a potential new hub for EP4 sustained activation (**Fig. 3L**). In contrast, we observed a decrease in signal for mGα protein recruitment to the ER consistent with the trafficking of EP4 away from this compartment (**Fig. 3M**).

These findings highlight the sustained activation of EP4 in multiple intracellular compartments and suggest broader functions beyond the plasma membrane.

### EP4 assembles Gα and β-arrestin signaling complexes in intracellular compartments

Conventional BRET is limited to studying the proximity between only two proteins. To surmount this limitation, NanoLuc Binary Technology BRET (NanoBiT-BRET, nbBRET) was used to simultaneously measure the proximity between tagged receptor, effector (m*G*α and β-arrestin) and localization marker (*34, 35*) (**Fig. 4A**). This split luciferase assay was developed by tagging EP4 at its C-terminus with the Natural Peptide fragment of NanoLuc (NatP, 13 residue NanoBiT fragment) and plasma membrane (CAAX), early endosomal (FYVE), late endosomal (Rab7a), recycling endosomal (Rab11a) or Golgi apparatus (Giantin) markers were tagged with the large NanoBiT fragment (LgBiT). Luminescence occurs from a complex between EP4-NatP and LgBiT-CAAX, LgBiT-FYVE, LgBiT-Rab7a, LgBiT-Rab11a or LgBiT-Giantin, which serves as an energy donor for fluorophore-tagged YFP-mGα or β-arrestin-YFP. In HEK293 cells, PGE_2_ (10 μM) increased nbBRET signal between EP4, mGαs, mGαi, β-arrestin-1 or β-arrestin-2 and LgBiT-CAAX, LgBiT-FYVE, LgBiT-Rab7a, LgBiT-Rab11a or LgBiT-Giantin, consistent with the recruitment of EP4, mGαs, mGαi, β-arrestin 1 and β-arrestin 2 to the plasma membrane, early, late and recycling endosomes and the cis-Golgi network. nbBRET signals were maintained for at least 20 min, suggesting assembly of stable EP4/mGαs/i or EP4/β-arrestin-1/2 complexes (**Fig. 4B-F**). PGE_2_ caused a concentration-dependent stimulation of complex assembly at the plasma membrane and in early endosomes (**Fig S1**), with potency values ranging between 14 nM and 288 nM. The antagonist for EP4, BGC20-1531 (3 µM), prevented PGE_2_ (100 nM)-stimulated assembly of EP4/mGαs, EP4/mGαi and EP4/β-arrestin-1 complexes in the plasma membrane and early endosomes (**Fig S2**).

**Figure 4:**
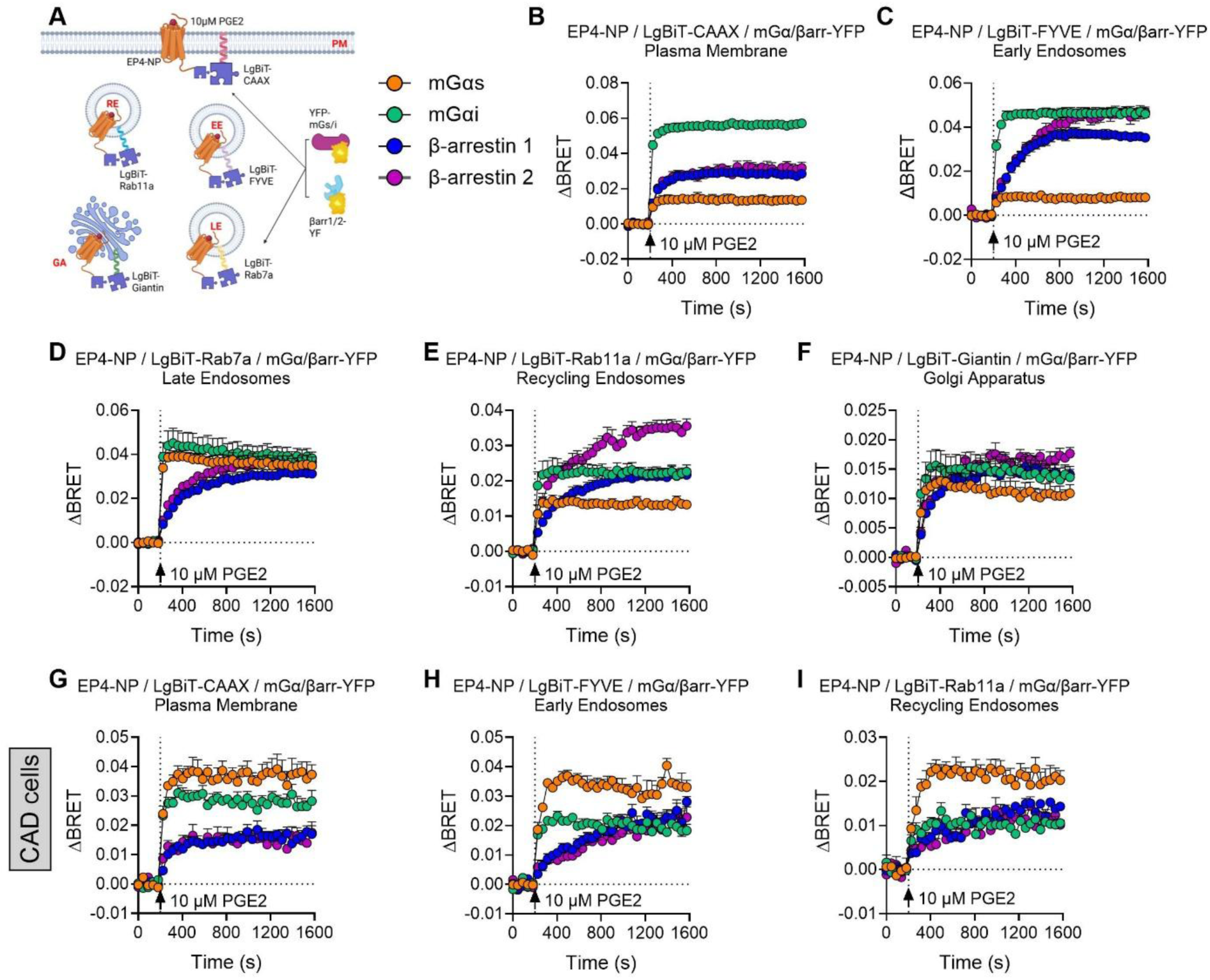
Assembly of EP4 signalosomes in HEK293 and in neuron-like CAD cells by NanoBiT-BRET (NbBRET). **A)** Illustration of NanoBiT-BRET (NbBRET) biosensors. NbBRET uses luciferase split into two fragments (Natural Peptide, NatP, and LgBiT) to detect BRET between receptor (EP4), effector (mGα or βARR) and proteins resident subcellular compartments. B-F). In HEK293 cells, PGE_2_ (10 µM) induced nbBRET between EP4-NatP, **B)** plasma membrane (LgBiT-CAAX), **C)** early endosomal (LgBiT-FYVE),**D)** late endosomal (LgBiT-Rab7a), **E)** recycling endosomal (LgBiT-Rab11a) or **F)** Golgi apparatus (LgBiT-Giantin) as well as YFP-mGαs, YFP-mGαi, β-arrestin1-YFP or β-arrestin2-YFP. G-i) In neuron-like CAD cells, PGE_2_ (10 µM) induced nbBRET between EP4-NatP, **G)** plasma membrane (LgBiT-CAAX), **H)** early endosomal (LgBiT-FYVE), or **I)** recycling endosomal (LgBiT-Rab11a) as well as YFP-mGαs, YFP-mGαi, β-arrestin1-YFP or β-arrestin2-YFP. Data are represented as the mean ± SEM (n=5).

To study signalosome assembly in a relevant cellular context, we also used nbBRET to study subcellular signalosomes of EP4 in CAD cells, a neuron-like cell line modified from the Cath.a catecholaminergic cell line obtained from a mouse tumor (*36*). In CAD cells, PGE_2_ (10 μM) also increased nbBRET signal between EP4, mGαs, mGαi, β-arrestin-1 or β-arrestin-2 and markers of the plasma membrane, early or recycling endosome. Effector recruitment was sustained for at least 20 min (**Fig. 4G-I**).

Considered together, these results support the hypothesis that PGE_2_ induces the recruitment of the EP4, mGαs, mGαi, β-arrestin-1 and β-arrestin-2 to endosomes and Golgi apparatus, where a multiprotein signalosome may transduce intracellular signals.

### EP4 activates G proteins in subcellular compartments

mGα proteins are valuable tools to detect receptor activation but do not inform on the presence or activation of Gα proteins within intracellular compartments. To assess the spatiotemporal dynamics of EP4-mediated Gα protein activation, we used the G protein Effector Membrane Translocation Assay (GEMTA) (*37*). GEMTA biosensors consist of subdomains of Gα protein effectors that selectively interact with activated Gα proteins and do not require Gα protein or receptor modification. To probe compartmentalized Gαi activation, we used ebBRET to monitor the EP4-mediated trafficking of Rap1GAP-RlucII, a Gαi effector, to intracellular compartments in HEK293 cells expressing different Gαi isoforms (Gαi1, Gαi2, Gαi3, Gαz, GαoA, GαoB) (**Fig. 5A**). This analysis enabled determination of the activity of Gαi in subcellular compartments. In parental HEK293 cells, EP4 activated Gαi1, Gαi2, Gαz and GαoB but not Gαi3 and GαoA at the plasma membrane. We observed rapid and sustained activation of Gαi1 and Gαi2, while Gαz exhibited slower activation and GαoB displayed a transient response (**Fig 5B**). We conducted the same experiment in βarrKO cells to investigate the impact of receptor endocytosis on G protein signaling. Unexpectedly, the absence of β-arrestins did not enhance Gαi activation at the plasma membrane, contrary to expectations considering their established role in GPCR desensitization (**Fig. 5C**).

**Figure 5:**
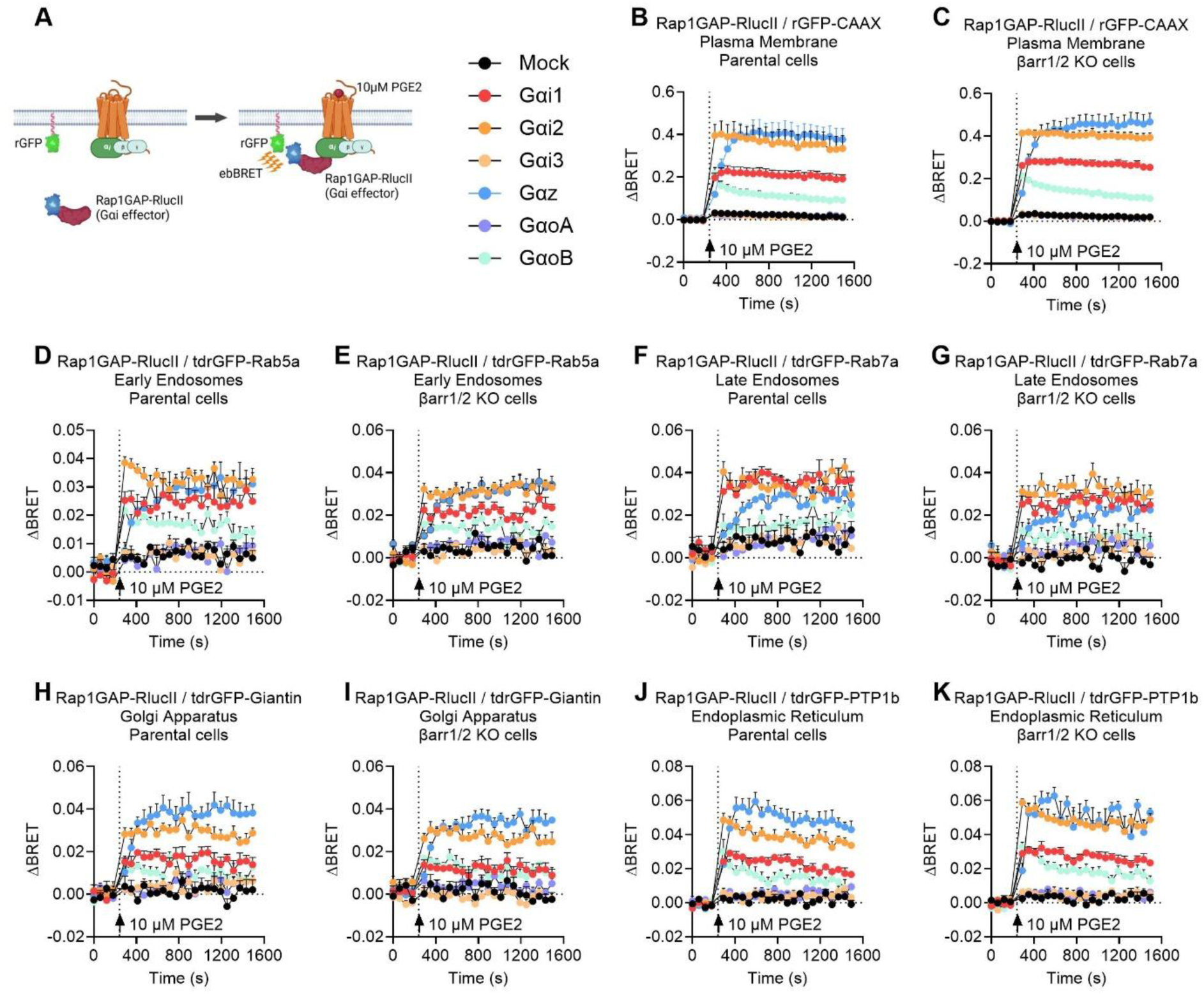
EP4-mediated compartmentalized Gαi activation. **A)** Illustration of the G protein Effector Membrane Translocation Assay (GEMTA) used to monitor Gi activation. Rap1GAP-RlucII is recruited by activated Gαi proteins in subcellular compartments. **B-K)** PGE_2_-induced (10μM) Gαi activation in different cellular compartments is monitored by transfecting parental HEK293 cells or β-arrestin1/2 KO cells with the Gαi effector Rap1GAP-RlucII, EP4, and ebBRET markers of **B-C)** the plasma membrane (rGFP-CAAX), **D-E)** early endosomes (tdrGFP-Rab5), **F-G)** late endosomes (tdrGFP-Rab7), **H-I)** Golgi apparatus (tdrGFP-Giantin), and **J-K)** endoplasmic reticulum (tdrGFP-PTP1b). Different Gαi subtypes (Gαi1, Gαi2, Gαi3, Gαz, GαoA and GαoB) are exogenously expressed to determine EP4 G protein selectivity. Data are represented as the mean ± SEM (n=4-6).

Next, we investigated EP4-mediated Gαi activation within endosomal compartments. In parental HEK293 cells, PGE_2_ stimulated sustained Gαi1, Gαi2, Gαz and GαoB activation in early endosomes (**Fig. 5D**). Surprisingly, β-arrestin 1/2 depletion did not inhibit Gαi activation as might be expected due to their role in EP4 endocytosis. Instead, Gαi1, Gαi2, Gαz and GαoB activation persisted, suggesting a distinct mechanism of Gαi activation in the endosomes that does not depend on β-arrestin-mediated EP4 internalization (**Fig. 5E**). Similarly, EP4 promoted Gαi1, Gαi2, Gαz and GαoB activation in late endosomes in parental and βarrKO cells, reinforcing the idea that EP4 internalization is not strictly required for endosomal Gαi activation (**Fig. 5F, G**). At the Golgi apparatus and ER, we detected Gαi1, Gαi2, Gαz and GαoB activation (**Fig. 5H, J**) that was maintained in absence of β-arrestin 1/2 (**Fig. 5I, K**) indicating that EP4-induced Gαi signaling persists beyond endosomal compartments. Gαi activation in recycling endosomes was also weakly detected in parental HEK293 cells and βarrKO cells (**Fig. S3**). Pertussis toxin (PTX) treatment blocked Gαi1, Gαi2 and GαoB activation, but not Gαz activation as this Gαi subtype is known to be insensitive to PTX (*38*) (**Fig. S4**).

Since EP4 robustly couples to Gαs, we measured PGE_2_-induced cAMP production using the Green Up cADDis cAMP sensor (*39, 40*) of total cellular cAMP production. We also used a plasma membrane-anchored cADDis sensor to specifically monitor cAMP production in this compartment (*41*). EP4 stimulation increased total cellular cAMP levels (**Fig. 6A, C**). Expression of the endocytosis inhibitor DynK44A significantly reduced total cAMP production presumably due to loss of Gαs signaling from intracellular compartments (**Fig. 6A, C**). However, when monitoring cAMP production specifically at the plasma membrane, we found that plasma membrane-localized cAMP production was not affected by DynK44A (**Fig. 6B, C**). This suggests that EP4 mediates cAMP production at the plasma membrane but also within intracellular compartments that can be reduced using DynK44A to inhibit receptor endocytosis and Gαs activation in intracellular organelles.

**Figure 6:**
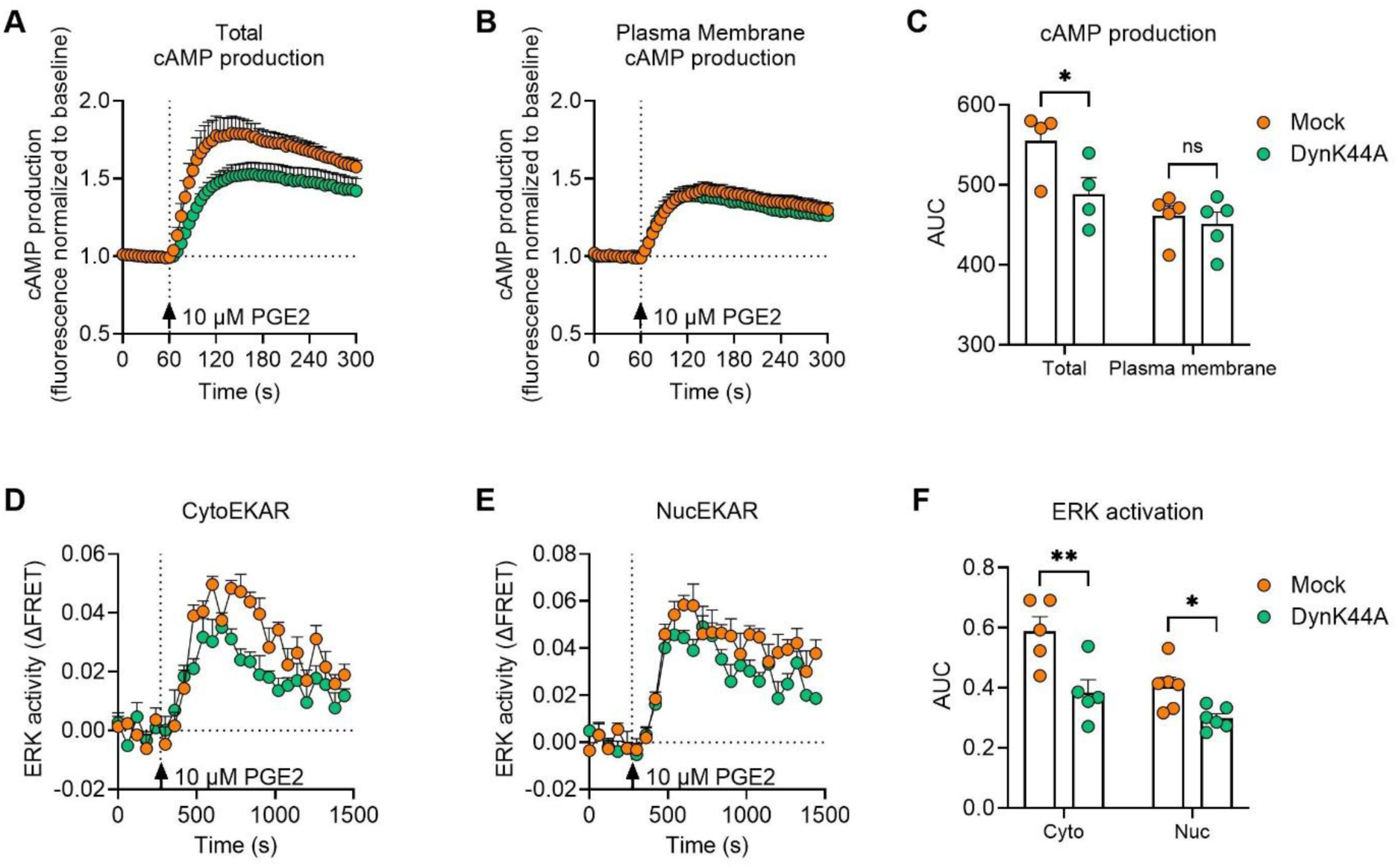
Spatiotemporal characterization of EP4-mediated intracellular Gαs signaling. PGE_2_-induced (10μM) **A)** total or **B)** plasma membrane localized cAMP production is measured in HEK293 cells transduced with respectively the cytosolic or plasma membrane-anchored Green Upward cADDIS cAMP reporter in presence or absence of the endocytosis inhibitor DynK44A. **C)** Quantification of cAMP production over time. Area under the curve (AUC) is used as a measure of response. ERK phosphorylation in the cytosol and nucleus was determined using the FRET sensors **D)** CytoEKAR and **E)** NucEKAR, respectively. **F)** Quantification of ERK activation over time. Area under the curve (AUC) is used as a measure of response. Data are represented as the mean ± SEM (n=4-6). Statistical significance of the differences was assessed using a two-way ANOVA followed by Holm-Šídák’s multiple comparison test (ns nonsignificant; *P ≤ 0.05; **P ≤ 0.01).

Given the established link between EP4-mediated cAMP signaling and downstream ERK activation, we examined the role of EP4 endocytosis on ERK activity using cytosolic (CytoEKAR) and nuclear (NucEKAR) FRET-based ERK biosensors (*42*). EP4 stimulation triggered robust ERK activation in both the cytoplasm and nucleus (**Fig. 6D, E**). However, in cells expressing DynK44A, ERK activation was significantly reduced in both compartments, with a more pronounced effect in the cytoplasm, indicating that EP4 endocytosis is required for efficient compartmentalized ERK signaling (**Fig. 6F**).

## Discussion

The dogma that GPCR signaling occurs exclusively at the plasma membrane and that β-arrestin-mediated desensitization and endocytosis terminate signaling has been refuted by studies that provide evidence that GPCRs can continue to signal from intracellular compartments by Gα protein and β-arrestin dependent and independent processes (*18–24*). Our previous work, along with others, showed that many GPCRs remain active after endocytosis and form signaling complexes with Gα proteins and β-arrestin within intracellular compartments. Our current results show that PGE_2_ stimulates EP4 receptor endocytosis and subsequent signaling in multiple intracellular compartments, and provides evidence that endocytosis is necessary for the sensitization of nociceptors and pain-like behavior. Using a combination of biosensors to measure EP4, Gα, β-arrestin, cAMP and ERK activity, our results indicate that EP4 signals from endosomal compartments, Golgi apparatus and endoplasmic reticulum, and suggest that EP4 endocytosis, while not required for Gαi activity, is important for persistent cAMP and ERK signaling.

EP4 is the principal mediator of PGE_2_-induced nociceptor sensitization and pain, primarily through direct activation of DRG nociceptor terminals *via* cAMP production and PKA signaling(*43–46*). This makes EP4 an attractive target for pain management. Several efforts have been made to develop EP4 receptor antagonists (*47, 48*), yet only one - grapiprant - has received FDA approval, and only for veterinary use in managing osteoarthritis-associated pain in dogs (*49*). Despite this promising precedent, no EP4 receptor antagonists have been approved for pain management in humans. This limited clinical translation may stem from the fact that current antagonists primarily act at the plasma membrane. Traditional drug discovery has focused on targeting GPCRs at the cell surface; however, plasma membrane signaling is often transient and tightly regulated, and may not fully account for sustained pathological signaling (*50*).

Clathrin- and dynamin-dependent endocytosis has been shown to mediate the internalization of multiple GPCRs in nociceptors and spinal neurons (*23, 24, 51*). Once internalized, endosomal signaling can activate kinases that contribute to sustained neuronal excitability required for nociception. In this study, we demonstrate that inhibition of endocytosis attenuates EP4-induced nociception and triggers both immediate and prolonged increases in nociceptor excitability, establishing a mechanistic link between EP4 endocytosis and pain.

Using ebBRET biosensors, we showed that EP4 receptor endocytosis is dynamin- and β-arrestin dependent. Expression of DynK44A, a dominant negative mutant harboring a mutation in its GTPase domain and impairs vesicle scission, effectively inhibited EP4 endocytosis (*52, 53*). Similarly, depletion of β-arrestin 1/2 in HEK293 cells abolished EP4 receptor endocytosis, consistent with the established role of β-arrestin in scaffolding components of the endocytosis machinery (*54, 55*). Interestingly, we also showed that the EP4 receptor traffics to the Golgi apparatus, a process that requires both dynamin and β-arrestin. This finding parallels the retrograde trafficking to the Golgi apparatus mediated by the retromer complex, as observed for the thyroid stimulating hormone receptor (TSHR) (*56*) and the parathyroid hormone receptor (PTHR) (*19*). PGE_2_ also induced the export of EP4 from the ER. ER exit is the first step in the trafficking of freshly synthesized GPCRs to the plasma membrane and our data indicate that dynamin is an essential component of this process, while β-arrestin is not.

We found that EP4 not only traffics to and from these intracellular compartments, but also remains active within them. Upon PGE_2_ stimulation, both β-arrestin isoforms were robustly recruited to the plasma membrane, and trafficked to the endosomes along with the receptor. This sustained association is consistent with the profile of a class B receptor, which is known to maintain β-arrestin engagement in contrast to class A receptors, which rapidly dissociate from β-arrestins following internalization (*17, 57*). β-arrestin was shown to scaffold MAPK signaling cascades from endosomes in association with internalized GPCRs (*58*). However, its role in the trafficking and signaling events from the Golgi apparatus or in the ER remains to be explored.

Our data also shows mGαs and mGαi recruitment by the EP4 receptor, consistent with the previously observed coupling to the Gαs and Gαi protein families (*37, 59*). It is noteworthy that Avet et al. reported additional coupling to the Gαq and Gα12 families, while Inoue et al. identified Gα12 as the primary G protein coupled to EP4. In contrast, Masuho et al. only detected Gs coupling (*60*). The co-trafficking of the EP4 receptor with mGα proteins and β-arrestin was confirmed by nbBRET that allows to monitor their simultaneous presence in a cellular compartment. This approach was also applied to a more physiologically relevant cell model in CAD cells, demonstrating the formation of PGE_2_-induced signaling complexes in intracellular compartments.

To further dissect the EP4-mediated spatiotemporal dynamics and selectivity of Gαi activation, we took advantage of the recently developed GEMTA, which does not require receptor or Gα modification and allows tracking activation of specific Gα subtypes (*37*). Although our data and that of others demonstrate robust coupling to the Gαi family, we observed a selective propensity of the EP4 receptor to activate its members. Specifically, EP4 activated Gαi1, Gαi2, Gαz and GαoB with varying maximal responses, while no activation was detected for Gαi3 or GαoA. We also observed subtype specific temporal characteristics, as Gαi1 and Gαi2 showed rapid and sustained activation, Gαz activated more slowly, and GαoB showed a transient response.

Unexpectedly, we found that deletion of β-arrestin 1/2 did not inhibit EP4-mediated Gαi activation in the endosomes, the Golgi apparatus or the endoplasmic reticulum. Given the established role of β-arrestin in GPCR endocytosis, and that our current data shows impaired EP4 internalization and trafficking in absence of β-arrestin, we expected that blocking EP4 endocytosis should abolish Gαi endosomal activation. On the contrary, Gαi activation persisted in early and late endosomes, Golgi apparatus and ER. This suggests a mechanism in which activated Gαi translocates to these compartments independently of EP4 receptor endocytosis or β-arrestin scaffolding. A recent study investigating the μ-opioid receptor (MOR) signaling reported similar findings, showing that MOR-induced Gαi endosomal activation does not require MOR internalization or the presence of activated MOR in the endosome membrane (*61*). Together, these results raise the possibility that Gαi, once activated at the plasma membrane, may remain in an active state long enough to translocate to intracellular compartments and initiate signaling events there.

Gαs endosomal signaling has been investigated for several GPCRs (*18–20, 62*). These studies demonstrate the requirement for receptor endocytosis and β-arrestin to enable Gαs trafficking and scaffolding endosomal signaling complex (*21, 63*). Our data show a decrease in EP4 and Gαs-mediated total cAMP production in the presence of an endocytosis inhibitor (DynK44A), suggesting that endocytosis is required for full cAMP accumulation, likely due to disruption of Gαs-mediated signaling from intracellular compartments. We also found that inhibition of receptor endocytosis impaired ERK activation in line with previous findings (*24, 64*). Given that ERK mediates PGE_2_-evoked sensitization of nociceptors (*65*), this mechanism may explain why EP4 endocytosis and consequent signaling underlie nociceptor sensitization and resultant pain-like behavior.

There are several limitations to our study. First, although our behavioral data link EP4 endocytosis to nociception in mice, the relevance of these mechanisms in human pain physiology remains to be determined. Second, we investigated EP4 trafficking and signaling in HEK293 and CAD cells using receptor overexpression systems, which may not accurately reflect the spatial organization and dynamics of endogenous receptors in native nociceptors. Third, while we identified selective Gαi isoforms activation, we cannot exclude that those differences in kinetics of activation could be due to variation in expression levels of each Gα protein. However, our findings align with those of Avet et al., who similarly reported different activation levels of Gαi1, Gαi2, Gαz and GαoB by EP4. Finally, we observed that Gαi activation in subcellular compartments was independent of receptor endocytosis but the precise mechanisms by which activated Gαi proteins reach and signal from intracellular sites remain speculative and warrant further investigation. Alternatively, these findings could point to the existence of intracellular receptor pools that can directly activate Gαi proteins independently of traditional trafficking pathways.

Together, our findings uncover distinct cellular mechanisms underlying compartmentalized intracellular G protein signaling by EP4 that contribute to pain sensitization. We reveal that Gαi-mediated signaling can occur independently of EP4 internalization, whereas Gαs-driven endosomal signaling depends on EP4 endocytosis. These results highlight the potential of targeting EP4 receptor trafficking and signaling as a strategy to achieve analgesia while avoiding the broad systemic side effects associated with conventional PG-inhibiting therapies.

## Materials and methods

### Animals

Male and female mice C57BL/6J (Charles River, RRID:IMSR_JAX:000664, 6–8 weeks old) were used. Mice were housed in a temperature- and humidity-controlled vivarium (12 h dark/light cycle, free access to food and water, 5 animals per cage). Behavioral studies followed Animal Research: Reporting of In Vivo Experiments (ARRIVE) guidelines. Animal experiments and sample collections were carried out according to the European Union (EU) guidelines for animal care procedures and Italian legislation (DLgs 26/2014) application of the EU Directive 2010/63/EU. Animal studies were approved by the Animal Ethics Committee of the University of Florence and the Italian Ministry of Health (permits no. 765/2019-PR, 288/2021-PR). Mice were randomly assigned to experimental groups; group size was based on previous similar studies. Investigators were blind to treatments.

### Treatment protocol

Mice received intraplantar (i.pl., 10 μl/site) injection of L-902,688 (5 nmol) or vehicle. Dyngo-4a (500 pmol), PitStop2 (500 pmol) were administered (10 μL, i.pl.) 30 min before the algogenic stimuli (i.pl.).

### Nocifensive behavior

Immediately after i.pl. injection, mice were placed inside a plexiglass chamber, and nocifensive behavior response was assessed for 15 min by measuring the time (sec) that the animal spent lifting, biting, licking, shaking the injected paw.

### Mechanical allodynia

The mechanical paw-withdrawal threshold was measured using von Frey filaments of increasing stiffness (0.02–2 g) applied to the plantar surface of the mouse hind paw, according to the up-and-down paradigm. The 50% mechanical paw withdrawal threshold (g) response was then calculated from the resulting scores. Mechanical paw-withdrawal threshold was measured at baseline and at different times following treatments.

#### Intrathecal administration of siRNA to mice

Mouse Dnm1 siRNA (L-043277-01-0005) and nontargeting control (CTR) siRNA (D-001810-10-05) were from Dharmacon (**Table S1**). The Dnm1 or CTR siRNA (1.25 mg) was mixed with PEI-based transfection reagent (in vivo-jetPEI, 201-50 G; Polyplus, Illkirch, France) in an 8:1 N:P ratio (PEI nitrogen to DNA phosphate ratio). The siRNA in vivo jetPEI mixture was administered to conscious mice by i.t. injection (L4-L5, 5 µL) 48 hours before DRG dissection for electrophysiology experiments. *Dnm1* mRNA expression in DRG was downregulated 48 hours after siRNA injection(*26*).

#### Patch clamp electrophysiology

Dnm1 or control siRNA was administered by i.t. injection as described above. After 48 hours, lumbar (L3-L5) DRG isolated from male C57BL/6J mice were collected in Ca2+/Mg2+-free HBSS and incubated in a papain solution (60 U) (Worthington) for 10 minutes at 37 °C. The papain solution was then replaced with collagenase II (4 mg/mL, Worthington) and dispase II (4.6 mg/mL, Sigma-Aldrich), mixed in Ca2+/Mg2+-free HBSS, and incubated for an additional 10 minutes at 37°C. The DRG was pelleted by centrifugation at low speed (800 rpm for 1 minute) and subsequently resuspended in DMEM/F12 medium (Invitrogen) supplemented with 10% FBS and penicillin-streptomycin. Neurons were mechanically dispersed through trituration with a fire-polished Pasteur pipette and plated onto 5 mm round glass coverslips (Warner Instruments) coated with poly-D-lysine (0.05 mg/ml, Invitrogen) and laminin (6 µg/ml, Gibco). Neurons were maintained at 37°C in a humidified atmosphere (95% air, 5% CO2). Patch clamp recordings were performed within 30 hours after neuronal plating. Perforated patch clamp recordings used a pipette solution containing amphotericin B (240 μg/ml, Thermo Scientific) in current clamp mode at room temperature. Patch electrodes, pulled from borosilicate glass on a micro-pipette puller, had a resistance of 3-4 MΩ. Recordings were carried out using an amplifier (Axopatch 200B, Molecular Devices) and an analog-to-digital converter (Digidata 1440A, Molecular Devices), controlled by a PC running pCLAMP 10 software. Data were filtered at 2 kHz and digitized at 10 kHz. Excitability changes, including rheobase and action potential firing frequency, were assessed using a ramp current protocol (100 pA, 1 second) in small-diameter DRG neurons (size: ≤ 25 µm). The rheobase was calculated as the minimum current required to trigger the first action potential during the stimulus ramp. Resting membrane potential was recorded in I=0 mode for each neuron before applying the ramp protocol. The recording chamber was continuously perfused with an external solution at a rate of 1 ml/min. The solutions had the following compositions (mM): pipette solution: K-gluconate 110, KCl 30, HEPES 10, MgCl2 1, CaCl2 2 (pH 7.2, adjusted with KOH; 290 mOsm); external solution: NaCl 143, KCl 5, HEPES 10, glucose 10, MgCl2 1, CaCl2 2 (pH 7.4, adjusted with NaOH; 305 mOsm).

### Cell lines

Parental HEK293 and β-arrestin 1/2 KO cells (provided by Dr Stephane Laporte, McGill University, Montreal, Quebec, Canada) were grown in complete Dulbecco’s modified Eagle’s medium (DMEM) supplemented with 10% fetal bovine serum (FBS) and 100 U/mL penicillin-streptomycin. CAD cells were grown in DMEM/F12 containing 8% FBS and 100 U/mL penicillin-streptomycin. Cells were maintained at 37°C in 5% CO2 and 95% O_2_.

### Enhanced bystander Bioluminescence Resonance Energy Transfer

To monitor receptor trafficking, parental HEK293 and β-arrestin 1/2 KO cells were transfected with EP4-Rluc8 (BRET donor) and rGFP-tagged intracellular compartment makers rGFP-CAAX, tdrGFP-Rab5a, tdrGFP-Rab7a, tdrGFP-Rab11a, tdrGFP-Giantin, or tdrGFP-PTP1b (BRET acceptors) with or without DynK44A expression.

To monitor mG protein and β-arrestin recruitment, HEK293 cells were transfected with EP4, Rluc8-tagged mG proteins (Rluc8-mG) or RlucII-tagged β-arrestin 1/2 (β-arrestin1/2-RlucII) and rGFP-tagged intracellular compartment makers rGFP-CAAX, tdrGFP-Rab5a, tdrGFP-Rab7a, tdrGFP-Rab11a, tdrGFP-Giantin, or tdrGFP-PTP1b (BRET acceptors).

To monitor compartmentalized G protein activation, parental HEK293 or β-arrestin 1/2 KO cells were transfected with EP4, Rap1GAP-RlucII (BRET donor), rGFP-tagged intracellular compartment makers rGFP-CAAX, tdrGFP-Rab5a, tdrGFP-Rab7a, tdrGFP-Rab11a, tdrGFP-Giantin, or tdrGFP-PTP1b (BRET acceptors) and different Gi subtypes as indicated. Cells were pretreated in the absence or presence of the Gαi/o inhibitor Pertussis toxin overnight (PTX 100 ng/mL).

Transfected cells were seeded (40000 cells/well) in white opaque 96-well plates (Greiner). 48 hours post-transfection, cells were washed with PBS and media was replaced by HBSS containing 10 mM HEPES at pH 7.4. Prolume Purple (2μM, Nanolight Technology) was added for 6 min before measuring baseline BRET signal for 3 minutes followed by cell stimulation with 10 μM PGE_2_ and BRET measurement for the indicated times. BRET signal was measured on a Synergy Neo2 Microplate reader (BioTek) using BRET2 filters (Donor filter: 400 nm; Acceptor filter: 510 nm). The BRET signal is calculated as the ratio of light emitted by the energy acceptor over the light emitted by the energy donor and ΔBRET is calculated by subtracting the BRET signal in the vehicle condition from the BRET signal in the stimulated condition.

### NanoBiT-BRET

cDNA encoding human EP4 were cloned with a NanoBiT tag (GVTGWRLCERILA, 13 amino acids) appended to the C-terminus via a flexible linker (LRPLGSSGGG). This tag uses the NanoBiT fragment from the natural peptide within the nanoluciferase. Localization markers CAAX, FYVE, Rab7a, Rab11a or Giantin tagged on the N-terminus with HA, a short linker (GGSG) and the LgBiT tag (159 amino acids). CAAX and FYVE were from A. Thomsen (New York University). HEK293 or CAD cells were plated in white opaque 96-well plates. The following day, cells were transiently transfected using PEI or Lipofectamine. HEK293 cells were transiently transfected with EP4-NatP (20 ng/well), LgBiT-CAAX, LgBiT-FYVE, LgBiT-Rab7a, LgBiT-Rab11a or LgBiT-Giantin (20 ng/well), as well as either YFP-mGs, YFP-mGi, β-arrestin1-YFP, or β-arrestin2-YFP (20 ng/well). HEK293 cells were transfected using PEI (Polysciences) at a 6:1 ratio of DNA to PEI. CAD cells were transfected with EP4-NatP (100 ng/well), LgBiT-CAAX, LgBiT-FYVE or LgBiT-Rab11a (100 ng/well) and either YFP-mGs, YFP-mGi, β-arrestin1-YFP, or β-arrestin2-YFP (100 ng/well). CAD cells were transfected using Lipofectamine 3000 (Invitrogen) in OptiMEM following manufacturer’s instructions. After 48 h transfection, cells were washed with assay buffer (HBSS containing 10 mM HEPES at pH 7.4). HEK293 cells were pre-incubated for 30 min at 37°C with vehicle or EP4 antagonist BGC20-1531 (3 µM). Furimazine substrate (10 µM) was added 10 min prior to NanoBiT experiments at 37°C. Following a baseline for 3 min, cells were challenged with PGE_2_ (0.1 nM - 30 µM) or vehicle. Luminescence emissions were then recorded every 45 sec for 20 min on a Synergy Neo2 Microplate reader using BRET1 filters (Donor filter: 480 nm; Acceptor filter: 530 nm). ΔBRET represents the BRET signal in the presence of agonist, minus the BRET signal over time in the presence of vehicle.

### cADDis cAMP sensor assay

HEK293 cells were transfected with EP4 (2 µg/10 cm dish), and pcDNA3.1 or DynK44A (1 µg/10 cm dish). After 24 h, cells were transferred to 96-well plates and transduced with Green Up cADDis sensor BacMam (#U0250G, Montana Molecular) or Green smAKAP-targeted cADDis sensor BacMam (#U0267G, Montana Molecular) according to the manufacturer’s instructions. 24 h post-transduction, cells were washed and incubated in Hanks’ buffered saline solution (HBSS, pH 7.4) containing 0.1% BSA. After 30 min (37°C), fluorescence was measured for 5 min using an automated epifluorescence microscope (Leica-Microsystems) at 5 s intervals. After 1 min baseline read, cells were stimulated with PGE_2_ (10 µM). Average of 1 min baseline was considered as initial fluorescence for each well. cAMP levels were expressed as a fractional change from baseline for each time point. Area under the curve (AUC) was determined for each replicate. 3 technical replicates were included for each condition and 5-6 biological replicates were performed for every experiment.

### FRET

HEK 293 Flp-In cells were transfected with EKAR FRET biosensors targeted to the nucleus or cytoplasm (2 µg/10 cm dish), EP4 (2 µg/10 cm dish), and pcDNA3.1 or DynK44A (1 µg/10 cm dish). After 24 h, cells were transferred to 96-well plates and serum-starved overnight. On the day of the assay, cells were washed and incubated in Hanks’ buffered saline solution (HBSS, pH 7.4) containing 0.1% BSA. After 30 min (37°C), FRET was measured at 60 s intervals (CLARIOstar, BMG Labtech). After 5 baseline reads, cells were stimulated with PGE_2_ (10 µM). ΔFRET represents the ratio (YFP/CFP), minus the mean from 5 initial reads and baseline-corrected to vehicle. Area under the curve (AUC) was determined for each replicate. 3 technical replicates were included for each condition and 5-6 biological replicates were performed for every experiment.

### Data analysis and statistics

Data are presented as mean ± SEM, determined using GraphPad Prism (8.0). Concentration-response data were fitted using non-linear regression analysis as log *vs.* response (three parameters) to determine EC_50_ values. Differences were assessed using paired or unpaired t-test for two comparisons, and 1- or 2-way ANOVA and Tukey’s, Dunnett’s or Šidák’s post-hoc test for multiple comparisons. *P<*0.05 was considered significant at the 95% confidence level.

## Acknowledgment

We thank Dr Michel Bouvier (University of Montreal, Montreal, Quebec, Canada) for providing the BRET sensor for this study.

## Author contributions

Conceptualization: BS, RT, FDL, RN, PG, and NWB. Investigation: BS, RT, PD, NB, YM, FDL, and RN. Supervision: BLS, DDJ, PG, and NWB. Writing-original draft: BS, RT, PD, YM, and NWB. Writing-review and editing: BS, RT, PD, FDL, RN, PG, and NWB.

## Data availability

Contact the corresponding author (N. W. Bunnett at) to obtain original data.

