## Supplementary Materials for "Endocytosis and Compartmentalized Intracellular Signaling of the Prostaglandin Receptor EP4 Mediate Pain"

Supplementary digital content

Figs. S1 to S4
Tables S1

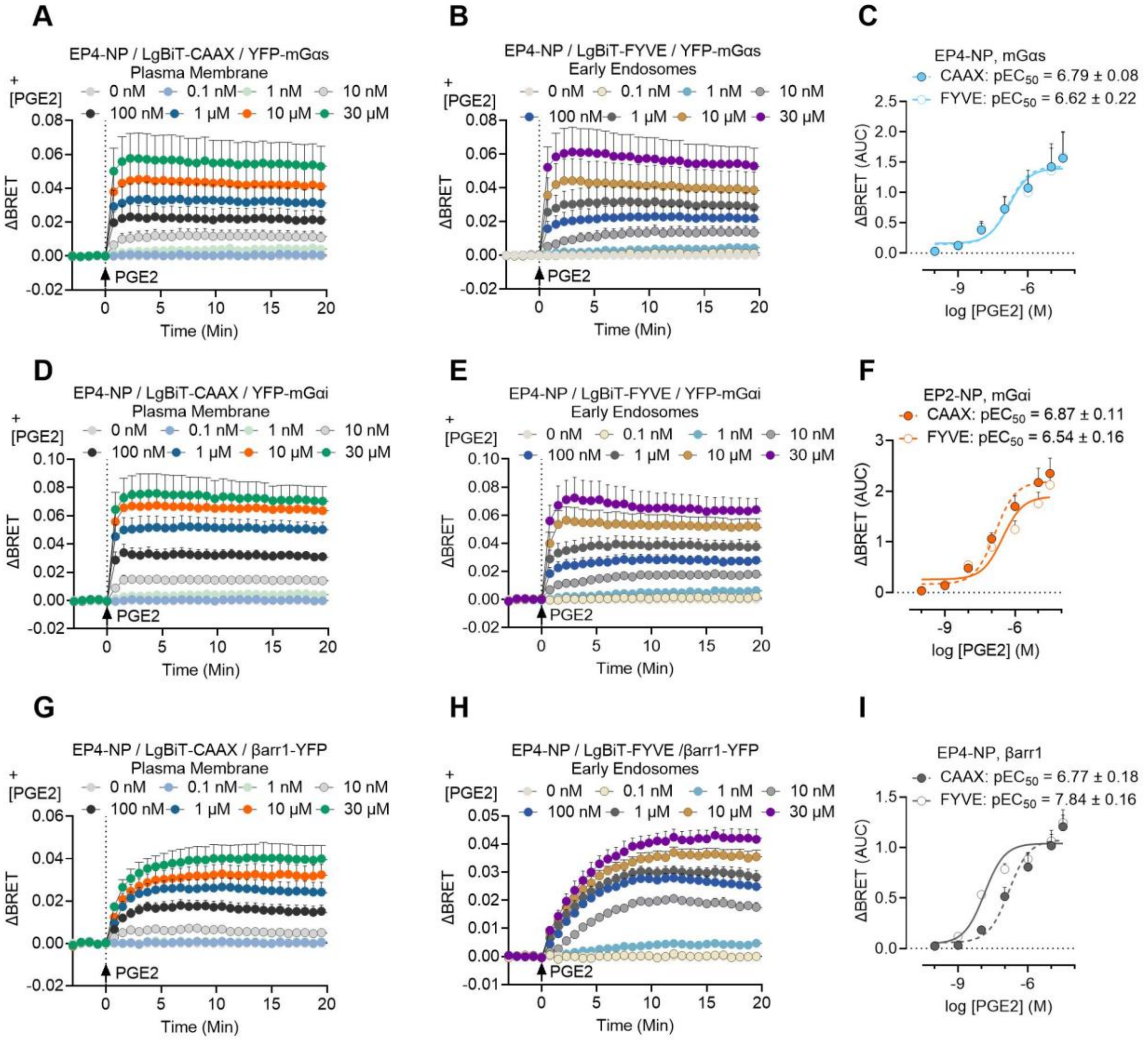


**Figure S1: Assembly of EP4 signalosomes in HEK293 cells.** Effects of graded concentrations of PGE2-induced nbBRET between EP4-NatP, LgBiT-CAAX (plasma membrane) or LgBiT-FYVE (early endosomes), as well as YFP-mGαs, YFP-mGαi or β-arrestin1-YFP. Data are represented as the mean ± SEM (n=5).

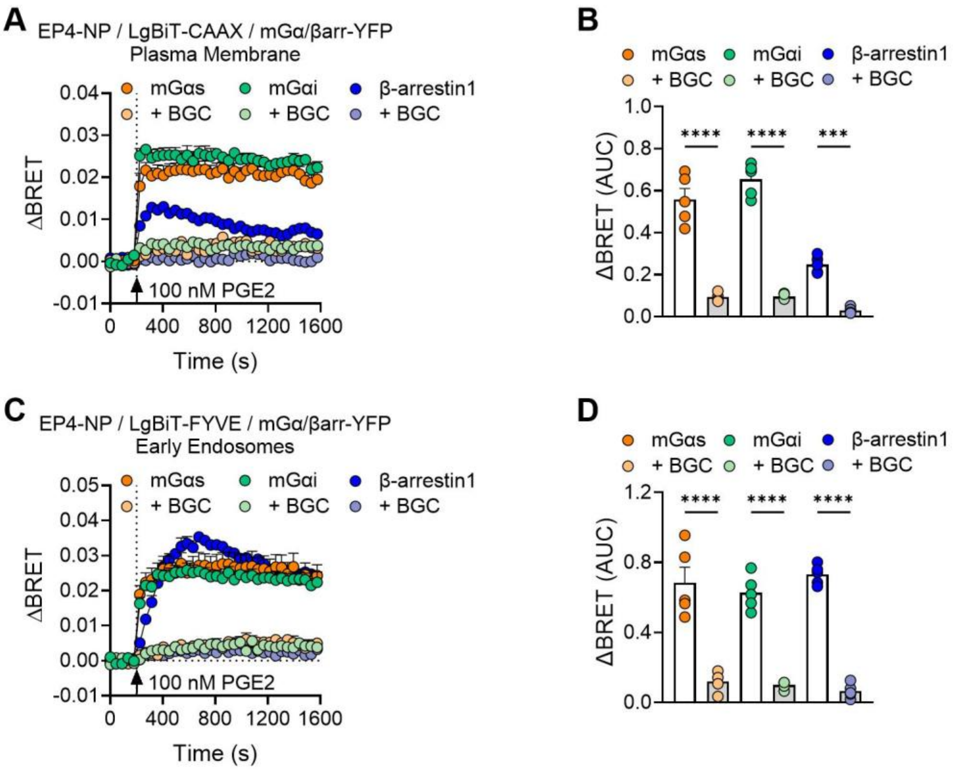


**Figure S2: EP4 antagonist inhibits EP4 signalosome assembly in HEK293 cells.** Effect of BGC20-1531, EP4 antagonist (BGC, 3 µM) on PGE2–induced recruitment of mGαs, mGαi and βARR1 to the **A, B)** plasma membrane (CAAX) or **C, D)** early endosome (FYVE) on NanoBiT-BRET (nbBRET) with natural peptide (NatP)–tagged EP4 in HEK293 cells. AUC, area under the curve. Data are represented as the mean ± SEM (n=5). 1-way ANOVA with Dunnett's test. ***P<0.001, ****P<0.0001 vs vehicle.

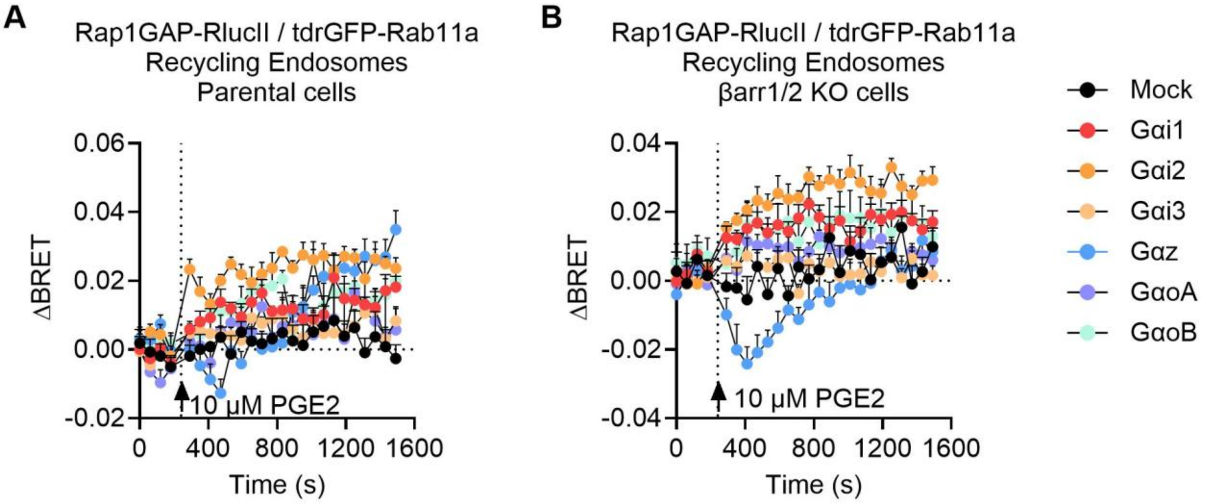


**Figure S3: EP4-mediated compartmentalized G protein activation in recycling endosomes.** PGE2-induced (10μM) Gαi activation in recycling endosomes is monitored by transfecting parental HEK293 cells or β-arrestin1/2 KO cells with the Gαi effector Rap1GAP-RlucII, EP4, and ebBRET marker tdrGFP-Rab11. Different Gαi subtypes (Gαi1, Gαi2, Gαi3, Gαz, GαoA and GαoB) are exogenously expressed to determine EP4 G protein selectivity. Data are represented as the mean ± SEM (n=5).

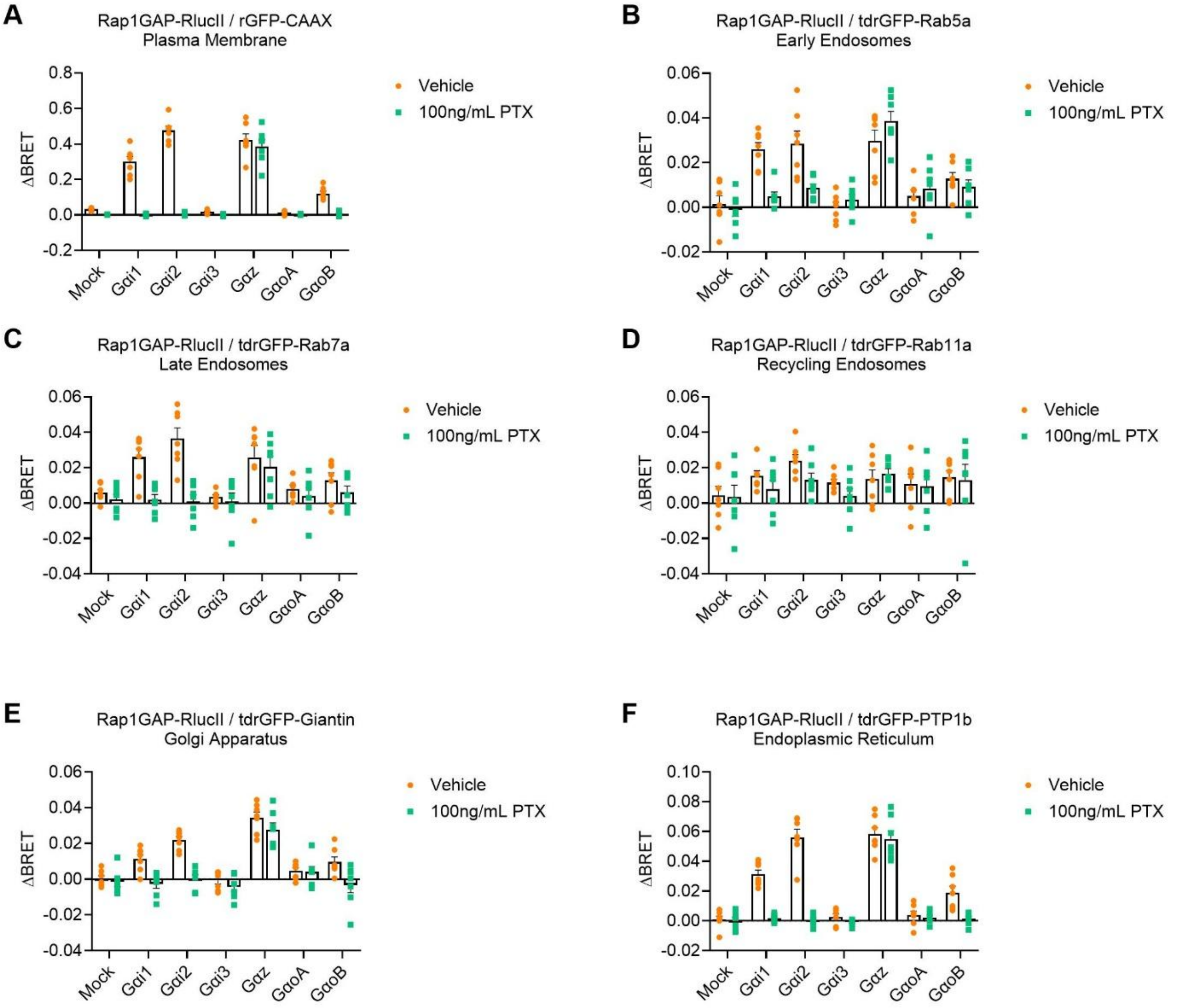


**Figure S4: Pertussis toxin inhibits EP4-mediated compartmentalized Gαi activation.** PGE2-induced (10µM) Gαi activation in different cellular compartments is monitored by transfecting parental HEK293 cells with the Gαi effector Rap1GAP-RlucII, EP4, and ebBRET markers of **A)** the plasma membrane (rGFP-CAAX), **B)** early endosomes (tdrGFP-Rab5a), **C)** late endosomes (tdrGFP-Rab7a), **D)** recycling endosomes (tdrGFP-Rab11a), **E)** Golgi apparatus (tdrGFP-Giantin), and **F)** endoplasmic reticulum (tdrGFP-PTP1b). Different Gαi subtypes (Gαi1, Gαi2, Gαi3, Gαz, GαoA and GαoB) are exogenously expressed to determine EP4 G protein selectivity. Cells were pretreated in the absence or presence of the Gαi/o inhibitor Pertussis toxin overnight (PTX 100 ng/mL). Data are represented as the mean ± SEM (n=7).

**Table S1.** Dnm1 and control siRNA sequences. m, mouse.

| Target | Dharmacon Sequence |
|---|---|
| *mDnm1* | GCGUGUACCCUGAGCGUGU, |
| | UGGUAUUGCUCCUGCGACA, |
| | GGGAGGAGAUGGAGCGAAU, |
| | GCUGAGACCGAUCGAGUCA. |
| *Control* | UGGUUUACAUGUCGACUAA, |
| | UGGUUUACAUGUUGUGUGA, |
| | UGGUUUACAUGUUUUCUGA, |
| | UGGUUUACAUGUUUUCCUA. |
